# Interactions between N-Terminal Modules in MPS1 Enable Spindle Checkpoint Silencing

**DOI:** 10.1101/438903

**Authors:** Spyridon T. Pachis, Yoshitaka Hiruma, Anastassis Perrakis, Geert J.P.L. Kops

## Abstract

Faithful chromosome segregation relies on the ability of the spindle assembly checkpoint (SAC) to delay anaphase onset until all chromosomes are attached to the mitotic spindle via their kinetochores. MPS1 kinase is recruited to unattached kinetochores to initiate SAC signaling, and is removed from kinetochores once stable microtubule attachments have been formed to allow normal mitotic progression. Here we show that a helical fragment within the kinetochore-targeting NTE module of MPS1 is required for interactions with kinetochores, and also forms intramolecular interactions with its adjacent TPR domain. Bypassing this NTE-TPR interaction results in high MPS1 levels at kinetochores due to loss of regulatory input into MPS1 localization, ineffecient MPS1 delocalization from kinetochores upon microtubule attachment, and SAC silencing defects. These results show that SAC responsiveness to attachments relies on regulated intramolecular interactions in MPS1 and highlight the sensitivity of mitosis to perturbations in the dynamics of the MSP1-NDC80-C interactions.

---

Genomic stability is a key feature of cellular homeostasis, and error-free chromosome segregation during mitosis is crucial for maintaining it ^1^. The spindle assembly checkpoint (SAC) safeguards this process by prohibiting cells from separating their duplicated chromosomes in anaphase until all of them are properly attached to microtubules of the mitotic spindle ^2,3^. The SAC is satisfied only when all chromosomes have made stable, bioriented attachments, a state in which each of the two sister chromatids is attached exclusively to microtubules emanating from opposing spindle poles. Any other attachment conformations are sensed and destabilized by the error correction machinery ^4^.

Unattached kinetochores elicit a SAC response by the hierarchical recruitment of SAC components including the BUBs (BUB1, BUB3, BUBR1) and the MADs (MAD1, MAD2). Eventually, this leads to the production of a diffusible anaphase inhibitor known as the mitotic checkpoint complex (MCC) ^5–8^. The MCC is responsible for blocking activation of the APC/C^CDC20^ complex whose function is to promote the transition to anaphase. SAC signaling is locally silenced when microtubules form stable attachments to a kinetochore, ultimately followed by full SAC satisfaction (MCC disassembly) when all chromosomes have achieved stable attachments ^3,9–11^.

Monopolar spindle 1 (MPS1) kinase is the major orchestrator of SAC signaling ^12^. It is recruited to unattached kinetochores where it autoactivates and subsequently phosphorylates its kinetochore substrates in order to recruit downstream SAC components and enable MCC production ^2,13–17^. MPS1 phosphorylates multiple residues on at least three proteins involved in the recruitment cascade (Knl1, Bub1, Mad1) ^18–23^ and may subsequently also directly impact MCC complex stability and its binding to the APC/C ^24^. Besides its role in the SAC, MPS1 is also involved in the regulation of chromosome biorientation ^25–28^.

MPS1 contains in its N-terminus an N-terminal extension (NTE) sequence module followed by a tetratricopeptide repeat (TRP) domain. Although the NTE provides the predominant localization signal, both modules are involved in MPS1’s ability to localize to kinetochores via direct binding to members of the NDC80 complex ^29–34^. The mitotic kinase Aurora B plays an important role in promoting MPS1 kinetochore localization ^27,35^, at least in part by alleviating an inhibitory effect that the TPR imposes on kinetochore binding via the NTE ^31^. MPS1 signaling at kinetochores diminishes upon microtubule binding due to competition between MPS1 and microtubules for binding to the NDC80 complex. Aided by high turnover of kinetochore MPS1 ^36,37^, this results in reduced ability of MPS1 molecules to re-bind kinetochores once they are occupied by microtubules ^32,33^. In this way SAC signaling is disrupted at its most upstream point - a mechanism that, alongside other ways of SAC silencing, eventually causes full SAC inactivation and anaphase onset ^38–48^.

The localization of MPS1 to kinetochores and regulation thereof are crucial for the SAC, yet much is still unknown about the mechanisms that dictate it. Here, we set out to examine the potential interplay between the different N-terminal regions in MPS1 and determine how they regulate its kinetochore levels. We report the presence of a helical fragment in the NTE of MPS1 that is important for its kinetochore localization, and we detect a direct intramolecular interaction between the NTE and TPR modules of MPS1. This interaction dampens MPS1 kinetochore binding which we show is important for efficient SAC silencing.

## RESULTS

### A short helical fragment in the NTE is important for MPS1 kinetochore localization

We have previously determined the crystallographic structure of Mps1^62-239^ allowing us to identify the TPR domain structure^31^. The last 40 residues in that structure were not visible in the electron density maps and are thus presumably flexible or disordered. To gain more insight into the structure and properties of MPS1’s N-terminal localization module, we performed NMR analysis on MPS1^1-239^, containing both the NTE module (which was not present in the crystallised protein), as well as the C-terminal extention (CTE) of the TPR domain (the last 40 residues that were not modeled). 87% of the residues in MPS1^1-239^ were successfully assigned to their corresponding NMR chemical shifts (**Fig S1a,b,c**). From the assignment it was evident that the NTE had chemical shifts characteristic of a flexible conformation. To further investigate that, we used the TALOS software ^49^ to predict secondary structure elements of the NTE based on the NMR spectra. TALOS confirmed that the NTE likely assumes a flexible conformation but also predicted (albeit with a low score) that residues 14-23 have the propensity to form an helix (**Fig 1a**). To examine the importance of this potential helical fragment, we designed a helix-disrupting point mutation in the NTE by substituting an Asparagine residue in the middle for a Proline residue (**Fig 1a,b**). Interestingly, MPS1 carrying the N18P mutation showed greatly reduced kinetochore levels in cells treated with nocodazole (**Fig 1c,d**). The low kinetochores levels of MPS1^N18P^ were very similar to those of MPS1^Δ60^, a mutant in which the entire NTE is removed (**Fig 1c,d)** ^31,50^. Combining the N18P substitution with a deletion of the TPR domain (MPS1^N18P-ΔTPR^) abolished MPS1 kinetochore binding to a similar extent as deleting the entire NTE-TPR module (MPS1^Δ200^) **(Fig 1c,d)** ^31^. We next measured mitotic delays in cells treated with nocodazole and a low dose of the MPS1 inhibitor Cpd-5 (25nM)^51^ to uncover potential subtle differences in SAC strength between the different cell lines. In agreement with compromised localization, cells expressing MPS1^N18P^ and MPS1^N18P-ΔTPR^ were impaired in maintaining a mitotic arrest when treated with nocodazole (**Fig 1e)**. Taken together, these data are consistent with a role for an N-terminal α-helix in enabling the interaction between the NTE and the NDC80-C.

**Figure 1.**
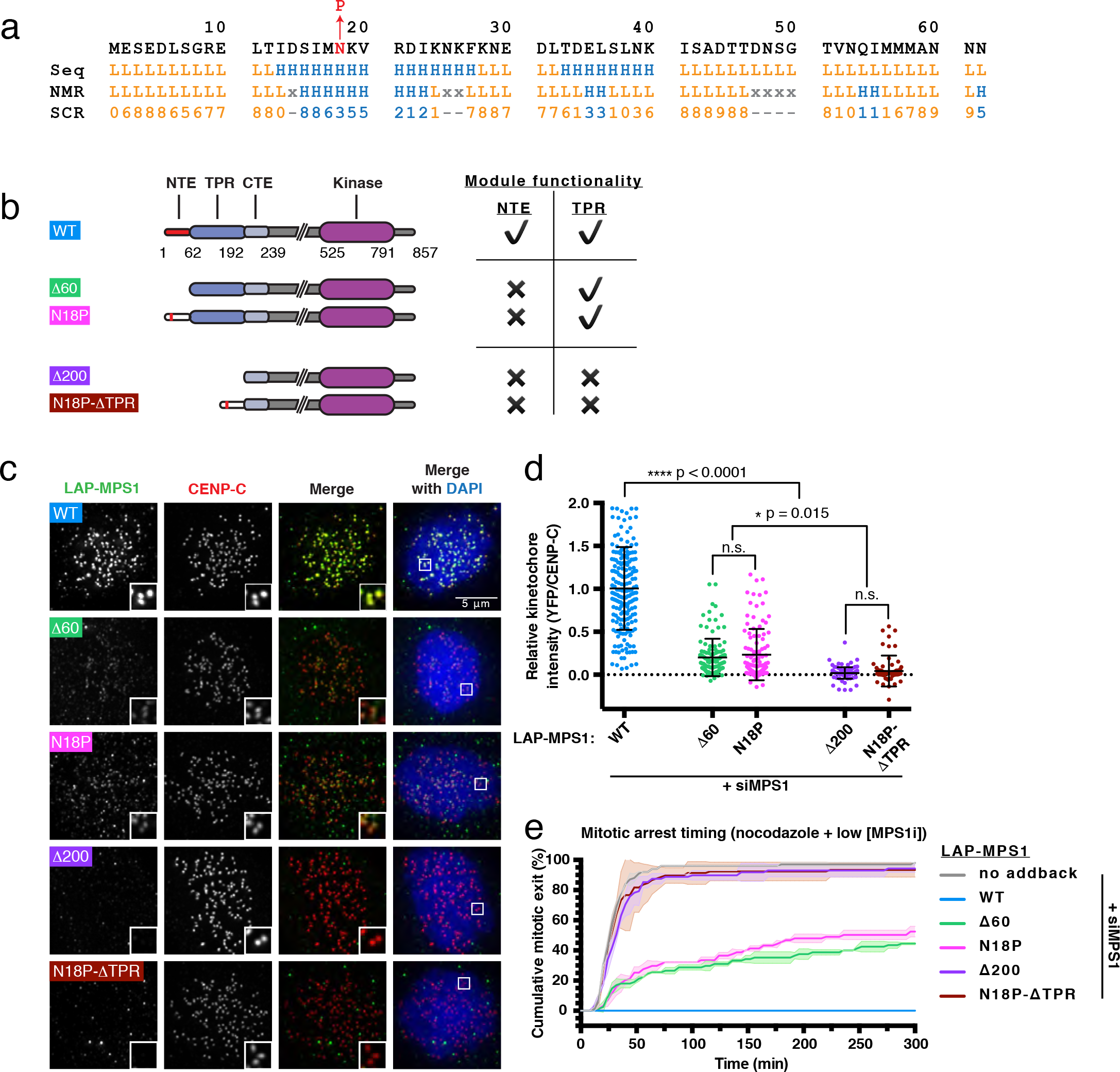
The NTE contains an α-helix that is important for MPS1 kinetochore localization. **(a)** Secondary structure prediction of the NTE. ‘Seq’ row shows the prediction based on the primary sequence, ‘NMR’ row shows the prediction based on the generated NMR spectra while ‘SCR’ is the confidence score per position for predictions based on NMR. **(b)** Schematic representation and functional classification of the different MPS1 variants. **(c and d)** Representative images **(c)** and quantification **(d)** of protein localization in nocodazole- and the MPS1 inhibitor Cpd-5 (25 nM)-treated HeLa FLP-In cells transfected with MPS1 siRNA and expressing the indicated LAP-MPS1 variants. The graph shows the mean kinetochore intensity (±s.d.) normalized to the values of MPS1^WT.^ Each dot represents one cell and all cells have been pooled from three independent experiments. Asterisks indicate significance (one-way ANOVA followed by Tukey’s test). **(e)** Time-lapse analysis of duration of mitotic arrest in nocodazole-and Cpd-5 (25 nM)-treated Hela Flp-In cells transfected with MPS1 siRNA and expressing the indicated LAP-MPS1 variants. Graph displays the mean values (±s.d.) from two independent experiments.

### The TPR domain and NTE of MPS1 interact

Our previous work suggested that regulation of MPS1 localization involves release of an inhibitory effect of the TPR domain on the NTE^31^. To examine if this could be via direct interactions between the two modules, we first assigned chemical shifts to MPS1^62-239^ (**Fig 2a**). We then compared the ^1^H, ^15^N HSQC spectra of MPS1^1-239^ and MPS1^62-239^ to measure chemical shift perturbations (CSPs) (**Fig. 2b**), which we mapped onto the sequence and crystal structure of the TPR domain (**Fig, 2c,d**). The largest CSPs, indicative of interaction sites with the NTE, were observed at the convex outer surface of the TPR, mostly at the two N-terminal α-helices (63−73 and 82−98) and considerably less so on the rest of the outer surface. Very few CSPs were measured at the concave inner surface of the TPR. To assess the contribution of electrostatic interactions to the interaction of the NTE with the TPR, we measured ^1^H, ^15^N HSQC spectra of the two MPS1 variants upon titration of KCl (**Fig S2a,d**). Whereas the chemical shifts of MPS1^62-239^ showed relatively small changes **(Fig S2a-c)**, notable spectral perturbations were observed for MPS1^1-239^ **(Fig S2d-h)**, suggesting that the NTE-TPR interactions are largely electrostatic. The N-terminal region of the TPR domain on the convex outer surface as well as the predicted helical residues in the NTE showed relatively large CSPs upon KCl titration, confirming that these two regions are involved in the interaction (**S3a-c**). These experiments suggest that the flexible NTE can interact with the proximal side of the convex outer surface of the TPR domain and this depends on electrostatic interactions.

**Figure 2.**
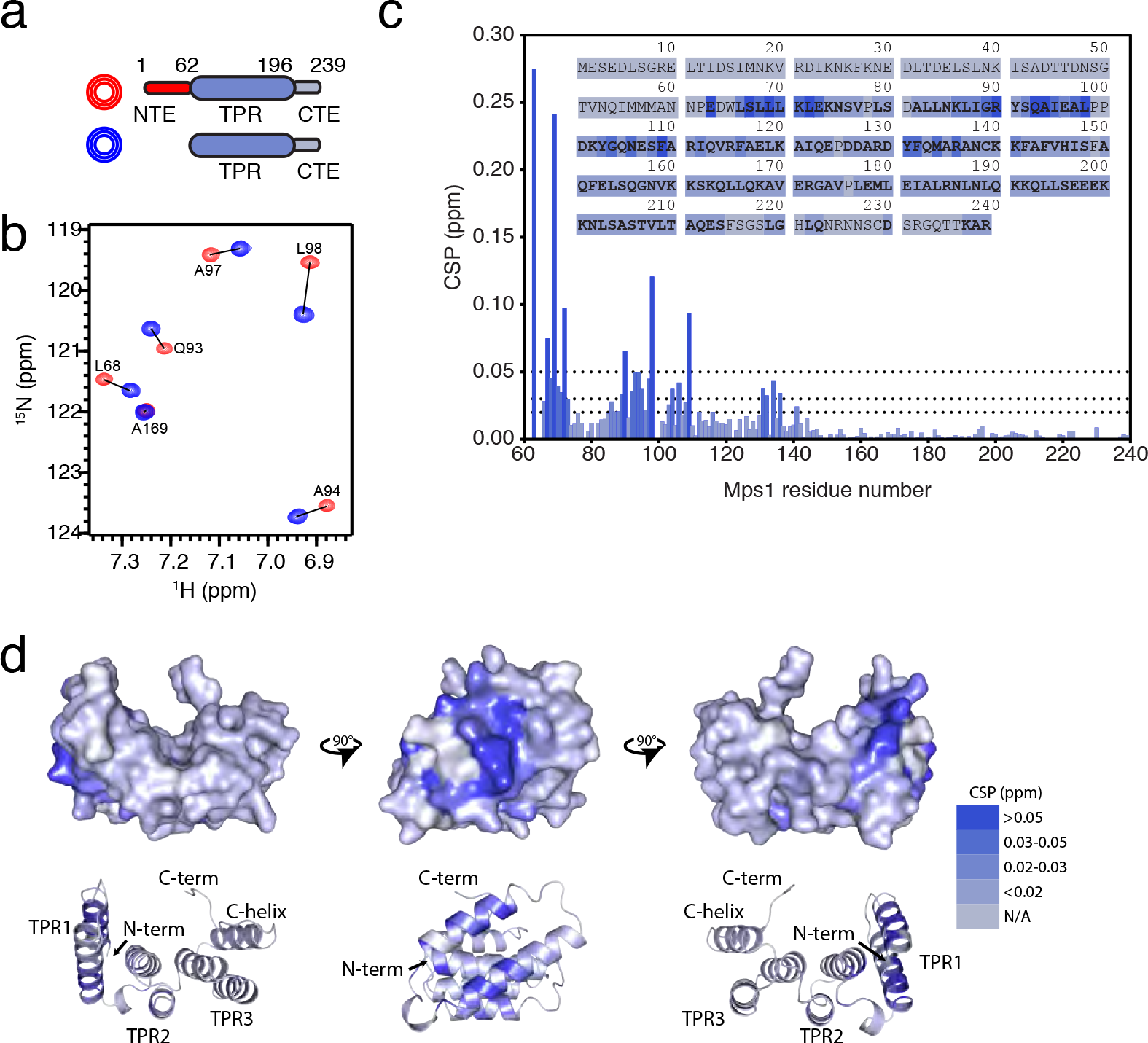
NMR of MPS1 N-terminal fragments uncovers NTE-TPR interactions. **(a)** Schematic representation of the variants of MPS1 that were used. **(b)** Examples of spectra showing residues in the TPR that are affected by the presence of the NTE (shifts from blue to red). **(c)** Display of the chemical shift perturbations (CSPs) per residue in the sequence of the TPR-CTE region, colour-coded based on the CSP magnitude. **(d)** Projection of the CSPs onto the crystal structure of the TPR domain (in 3 orientations) following the same colour-coding as in (c).

### Bypassing the NTE-TPR interaction leads to constitutively elevated levels of MPS1 on kinetochores

We next wished to examine the functional relevance of having an NTE-TPR interaction that can be regulated. We reasoned that if Aurora B activity controls MPS1 kinetochore localization by regulating this interaction as suggested by our prior work ^31^, then artificially circumventing it should render it insensitive to Aurora B. To achieve this, we created MPS1^2xNTE^, a version of MPS1 that contains an additional NTE fused to its N-terminus (**Fig 3a**). As predicted, MPS1^2xNTE^ kinetochore levels were largely insensitive to inhibition of Aurora B kinase activity (**Fig 3b,c**). Suprisingly, however, they were roughly two times higher than those of MPS1^WT^, and were additionally insensitive to inhibition of MPS1 activity, in contrast to MPS1^WT^ (**Fig 3b,c**). Enhanced levels of MPS1^2xNTE^ on kinetochores and insensitivity to MPS1 inhibition could not be explained by compromised kinase activity of MPS1^2xNTE 37,52-54^, as cells expressing this mutant were able to efficiently support the SAC in cells treated with nocodazole and a low dose of Cpd-5 (**Fig 3d**). These results indicate that the interaction between the NTE and the TPR domain of MPS1 is important for the regulation of MPS1 kinetochore levels and that Aurora B affects them, at least in part, through the regulation of that interaction.

**Figure 3.**
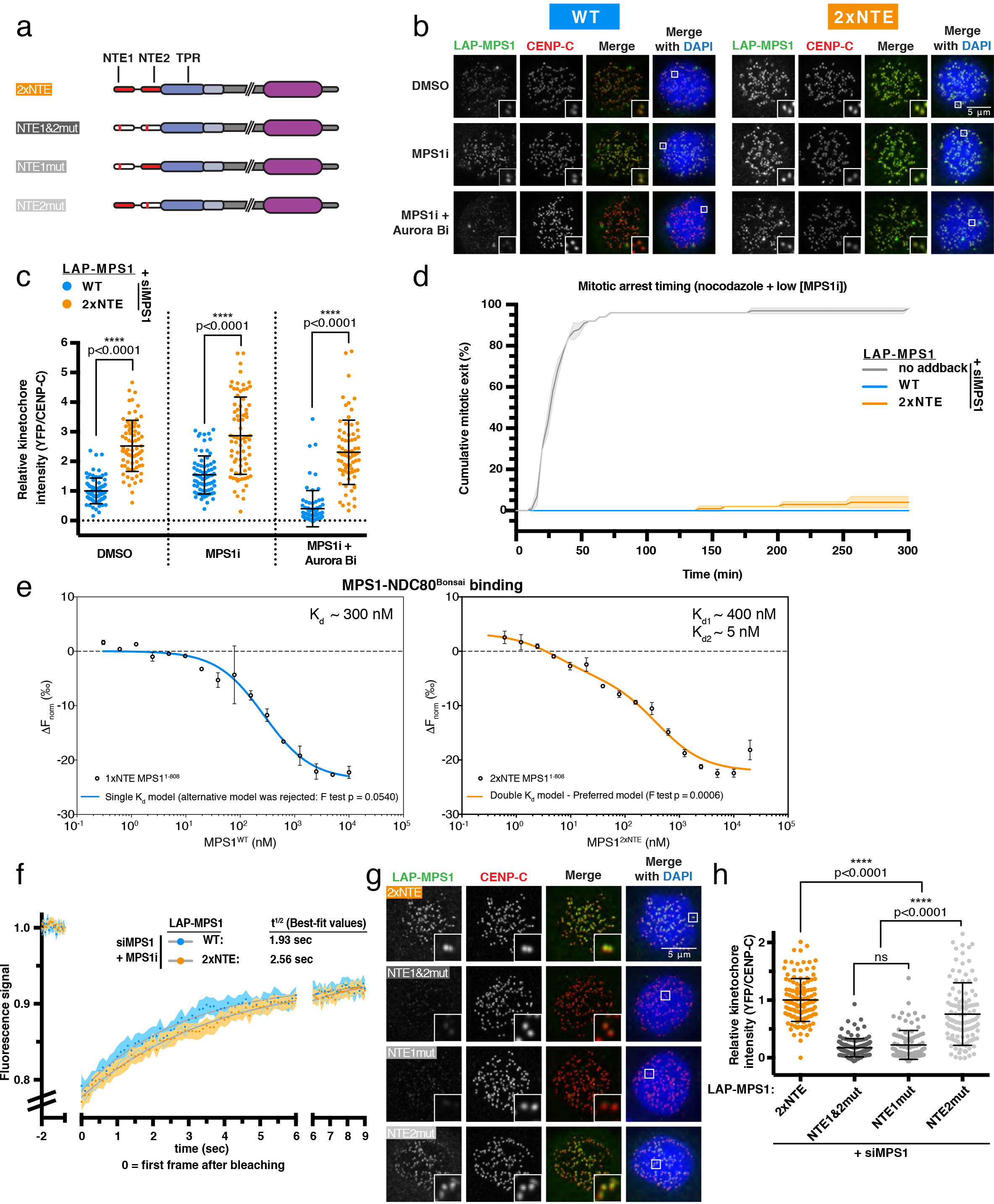
Bypassing the NTE-TPR interaction abolishes kinase regulation of MPS1 kinetochore localization. **(a)** Schematic representation of the MPS1 variants used. **(b and c**) Representative images **(b)** and quantification **(c)** of protein levels at kinetochores in HeLa Flp-In cells transfected with MPS1 siRNA, expressing the indicated LAP-MPS1 variants and treated with nocodazole and the indicated inhibitors. The graph shows the mean kinetochore intensity (±s.d.) normalized to the values of MPS1^WT^ in DMSO. Each dot represents one cell and cells have been pooled from four independent experiments. Asterisks indicate significance (one-way ANOVA followed by Tukey’s test). **(d)** Time-lapse analysis of duration of mitotic arrest in nocodazole- and Cpd-5 (25 nM)-treated Hela Flp-In cells transfected with MPS1 siRNA and expressing the indicated LAP-MPS1 variants. Graph displays the mean values (±s.d.) from three independent experiments. **(e)** MST binding graph generated by titrating MPS1^WT^ (left panel) or MPS1^2xNTE^ (right panel) to 50nM of NDC80^Bonsai^. One- or two-site binding curves were fitted and F test was performed to select a preferred model. **(f)** Quantification of FRAP performed on individual kinetochore pairs of nocodazole- and Cpd-5 (250 nM)-treated HeLa Flp-In cells transfected with MPS1 siRNA and expressing the indicated LAP-MPS1 variants. The graph displays the mean fluorescence intensity (±SEM) from two independent experiments **(g and h)** Representative images **(g)** and quantification **(h)** of protein levels on kinetochores in nocodazole- and Cpd-5 (250 nM)-treated HeLa Flp-In cells transfected with MPS1 siRNA and expressing the indicated LAP-MPS1 variants. The graph shows the mean kinetochore intensity (±s.d.) normalized to the values of MPS1^2xNTE^. Each dot represents one cell and cells have been pooled from 5 independent experiments. Asterisks indicate significance (one-way ANOVA followed by Tukey’s test).

### The NTE-TPR interaction promotes MPS1 release from kinetochores

We next tried to understand the nature of the increased kinetochore localization of MPS1^2xNTE^ and its insensitivity to kinase regulation. Kinetochore levels of both MPS1^WT^ and MPS1^2xNTE^ in cells treated with nocodazole were diminished upon knockdown of endogenous HEC1 by siRNA, confirming that MPS1^2xNTE^ kinetochore binding is dependent on the NDC80-C (**Fig S4a,b**). To examine whether the elevated kinetochore levels of MPS1^2xNTE^ are a result of a higher binding affinity between MPS1 and HEC1, we purified full-length recombinant MPS1 variants from insect cells and titrated them to a fixed concentration of NDC80-C^Bonsai 55^ in microscale thermophoresis (MST) experiments. MPS1^WT^ binding to NDC80-C^Bonsai^ fit a model consistent with a single binding event with a K_d_ of ^~^300 nM (**Fig 3e**, left panel). The binding of MPS1^2xNTE^ to NDC80-C^Bonsai^, however, is best described by a model with two binding events: one with a K_d_ of ~400 nM and one of ~5 nM (**Fig 3e**, right panel). Similar results were obtained when using shorter versions of MPS1 (1-377) (**Fig S5a**). The appearance of the higher affinity binding event is likely due to the added, unregulated NTE module.

We next performed FRAP analysis of fluorescent MPS1 variants at single kinetochore pairs of cells that were arrested in nocodazole to determine if differences in residence times contribute to the differences in kinetochore levels. We did so in the presence of MPS1 inhibitor (Cpd-5) to exclude activity-related effects on its residence time ^37^. Consistent with the presence of a higher affinity binding site on MPS1^2xNTE^ for the NDC80-C, MPS1^2xNTE^ displayed reduced turnover on kinetochores compared to MPS1^WT^ with best-fit recovery halftimes at 2.56 sec and 1.93 sec respectively (95% confidence intervals of halftime: WT 1.65-2.34 sec vs 2xNTE 2.19-3.0 sec) (**Fig 3f**).

To determine the contribution of each NTE to kinetochore binding, we first introduced the N18P mutation to disrupt the helical fragments present in both NTEs (MPS1^NTE1&2mut^) (**Fig 3a**). This severely compromised the ability of MPS1 to localize to kinetochores (**Fig 3g,h**). Strikingly, whereas mutation of only the TPR-proximal NTE (MPS1^NTE2mut)^ left localization of MPS1 largely unaffected, mutation of the apical NTE (MPS1^NTE1mut^) strongly reduced localization to levels similar to MPS1^NTE1&2mut.^ (**Fig 3g,h**). This observation argues that the increased levels of MPS1^2xNTE^ on kinetochores as well as the higher binding affinity in the MST experiments are mediated predominantly by the apical N-terminal NTE. Our data show that bypassing the NTE-TPR interaction removes kinase regulatory input into MPS1 localization and creates a higher affinity binding site for the NDC80-C which in turn leads to higher residence time of MPS1 on kinetochores.

### Bypassing the NTE-TPR interaction causes SAC silencing defects

Removal of active MPS1 from kinetochores is a crucial step for silencing the SAC in metaphase ^32,33,37^. Since kinetochore release of MPS1^2xNTE^ is compromised, we next examined whether removal of MPS1 from metaphase kinetochores was affected, by arresting cells with the proteasome inhibitor MG132. In contrast to MPS1^WT^, MPS1^2xNTE^ displayed significant retention on metaphase kinetochores, at levels approximately half of those observed in nocodazole-treated cells expressing the mutant (**Fig 4a,b**). Time-lapse imaging of cells going through a round of unperturbed mitosis revealed that MPS1 retention on metaphase kinetochores was accompanied by a mitotic arrest. Whereas around 95% of MPS1^WT^-expressing cells exited mitosis within 80 minutes from nuclear envelope breakdown (NEB), only 40% of MPS1^2xNTE^-expressing cells managed to complete mitosis even within 400 minutes (**Fig 4c**). The arrest was dependent on MPS1 activity (**Fig 4c**) and, consistently, was marked by elevated levels of KNL1-pT180, MAD1 and BUB1 on metaphase kinetochores. Of note, while KNL1-pT180 and BUB1 were present at roughly 70% of the levels observed in nocodazole-treated cells, MAD1 was retained at only 15% (**Fig 4d,e**, **S6a,b**).

**Figure 4.**
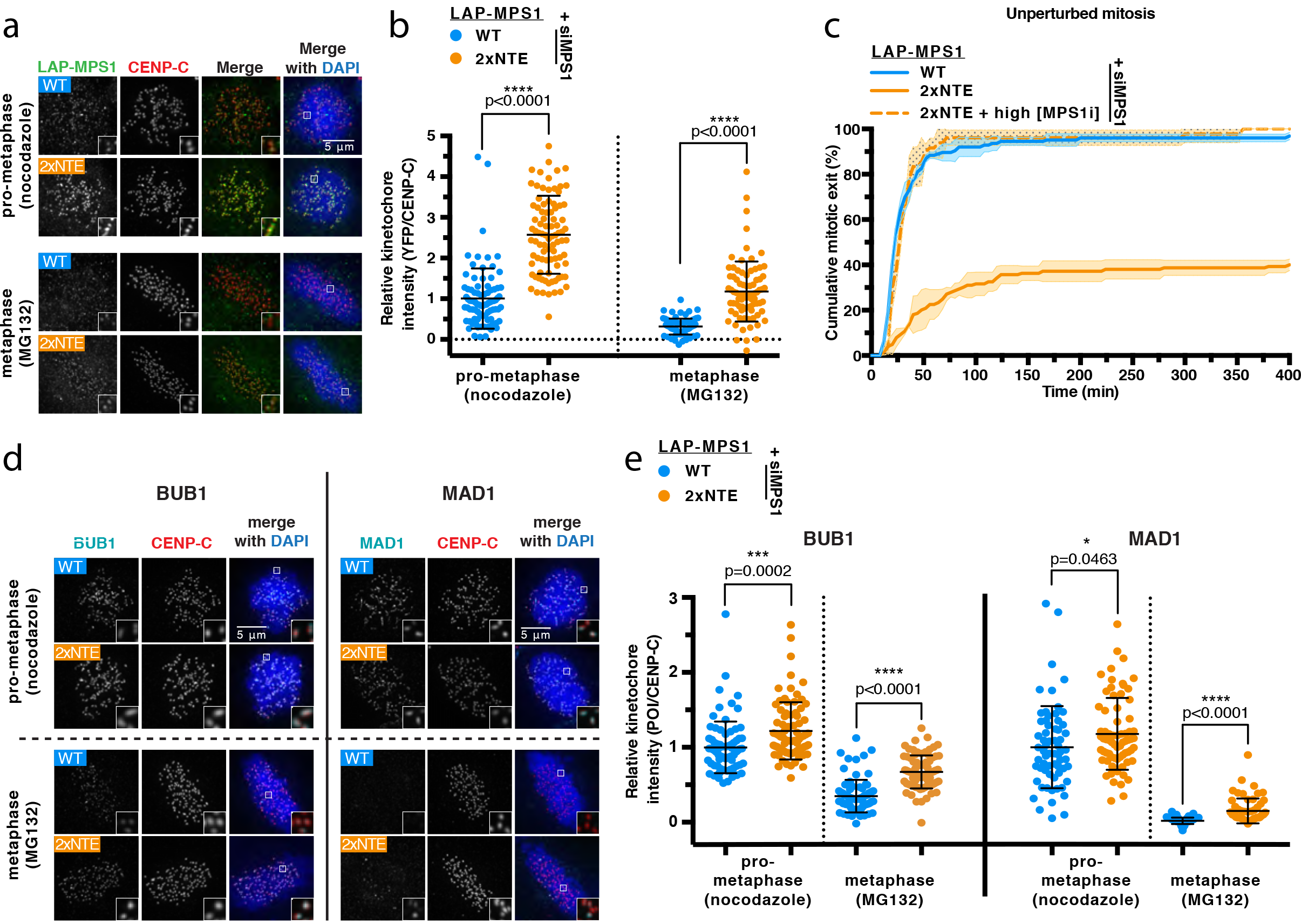
Bypassing the NTE-TPR interaction delays SAC silencing. **(a and b)** Representative images **(a)** and quantification **(b)** of nocodazole- or MG132-treated HeLa Flp-In cells transfected with MPS1 siRNA and expressing the indicated LAP-MPS1 variants. The graph displays the mean kineto-chore intensity (±s.d.) normalized to the levels of MPS1^WT^ in prometaphase. Each dot represents one cell and cells were pooled from 4 independent experiments. Asterisks indicate significance (student’s t test between the cell lines for each condition). **(c)** Time-lapse analysis of the duration of DMSO- or the MPS1 inhibitor Cpd-5 (250 nM)-treated HeLa Flp-In cells transfected with MPS1 siRNA and expressing the indicated LAP-MPS1 variants. Graph displays the mean values (±s.d.) from three independent experiments. **(d and e)** Representative images **(d)** and quantification **(e)** of the kinetochore levels of the indicated proteins in nocodazole- or MG132-treated HeLa Flp-In cells transfected with MPS1 siRNA and expressing the indicated LAP-MPS1 variants. The graph displays the mean kinetochore intensity (±s.d.) normalized to the levels of each protein in prometaphase MPS1^WT^-expressing cells. For each protein examined, cells were pooled from three independent experiments and each dot represents one cell. Asterisks indicate significance (studen’s t test between the different cell lines for each protein and condition).

We next examined whether weakened microtubule attachments due to the presence of elevated MPS1^2xNTE^ on metaphase kinetochores contributed to the mitotic arrest of the cells expressing this variant. Interestingly, kinetochore-microtubule attachments were unaffected, as indicated by four observations. First, the time from NEB to metaphase was indistinguishable between cells expressing MPS1^WT^ and MPS1^2xNTE^ (**Fig 5a,b**). Second, the time from metaphase to cohesion loss by cohesion fatigue ^56^ in MG132-treated cells was likewise similar (**Fig 5c**). Third, total levels of cold-stable tubulin of the mitotic spindle as well as individual k-fiber intensities were not substantially altered between cells expressing the two MPS1 variants (**Fig. 5d,e,f**) and no obvious correlation existed between the level of MPS1^2xNTE^ on single kinetochores and the k-fiber intensity (**Fig S6c**). Lastly, the levels of astrin, a marker of stable end-on attachments ^57^, were not reduced on metaphase kinetochores of cells expressing MPS1^2xNTE^, even though MPS1^2xNTE^ was still present there at high levels (**Fig 5g,h**). Taken together, these data show that bypassing the NTE-TPR interaction compromises the ability to silence the SAC by affecting the efficiency of MPS1 delocalization in metaphase without affecting microtubule attachments.

**Figure 5.**
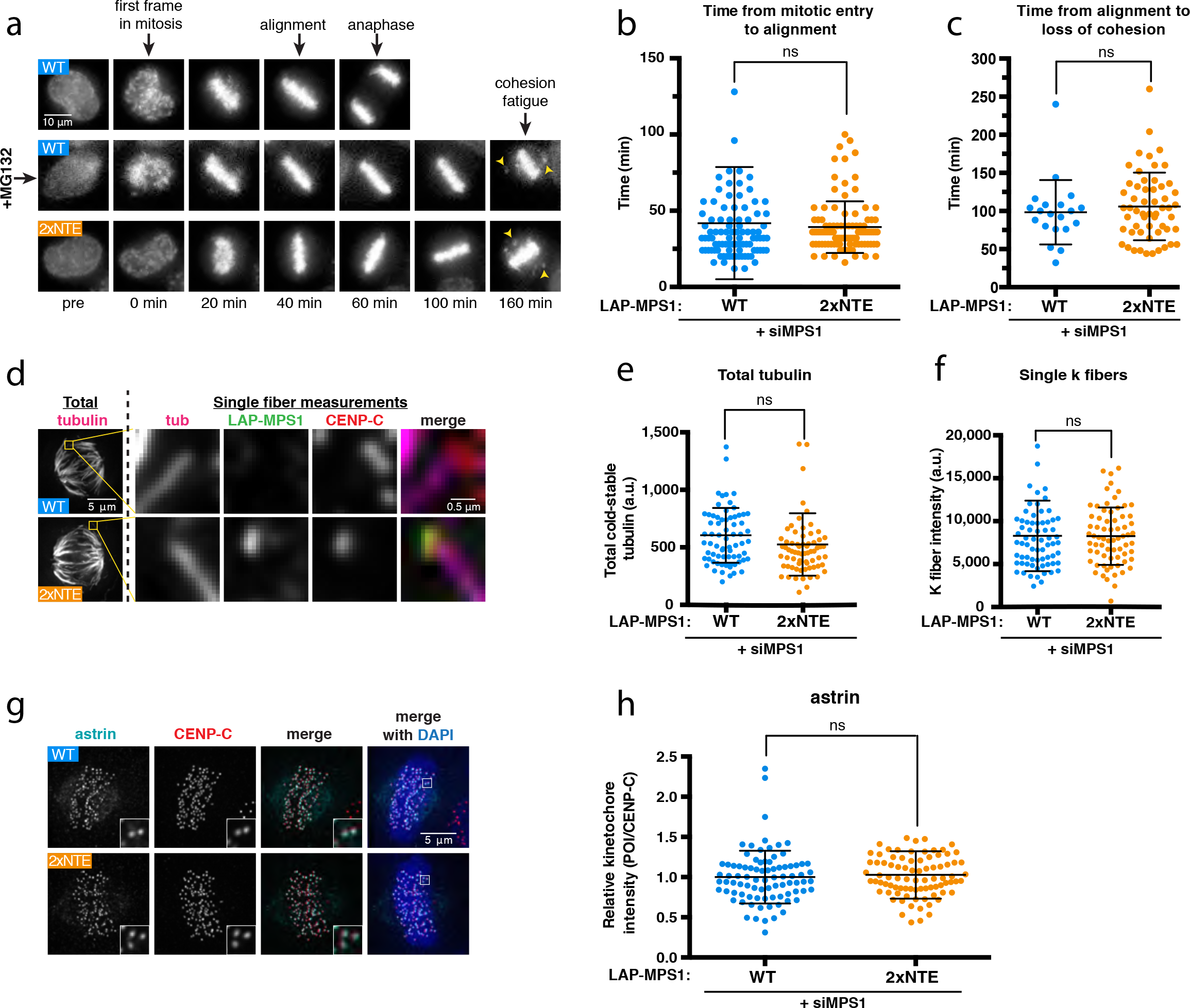
High levels of MPS1 on kinetochores in metaphase do not perturb microtubule attachments. **(a-c)** Representative stills **(a)** and quantifications **(b,c)** of timelapse movies of HeLa Flp-In cells transfected with MPS1 siRNA and expressing the indicated LAP-MPS1 variants. Cells were incubated with SiR-DNA to visualize the chromatin, and treated with either DMSO (2xNTE) or MG132 (WT). Graphs in (b,c) display the absolute time in minutes (±s.d.). Each dot represents a cell and cells were pooled from three (b) or two (c) independent experiments. Students t-test showed no significant differences. **(d-f)** Representative images **(d)** and quantifications **(e,f)** of cold-stable microtubules and levels of the indicated proteins on attached kinetochores in HeLa Flp-In cells transfected with MPS1 siRNA and expressing the indicated LAP-MPS1 variants. Graphs in (e,f) show quantification (a.u. ±s.d.) of the total cold-stable tubulin levels **(e)** or of intensity of individual k-fibers in each cell line **(f)**. Each dot represents a cell and cells were pooled from three independent experiments. Students t-test showed no significant differences. **(g and h)** Representative images **(g)** and quantification **(h)** of the kinetochore levels of astrin in MG132-treated HeLa Flp-In cells transfected with MPS1 siRNA and expressing the indicated LAP-MPS1 variants. The graph displays the mean kinetochore intensity (±s.d.) normalized to the levels of astrin in MPS1^WT^-expressing cells. Cells were pooled from two independent experiments and each dot represents one cell. Student’s t test was performed to determine significant changes.

## DISCUSSION

With this work, we uncover a short helical fragment in the NTE of MPS1, that is important for interaction with its kinetochore receptor. We also report a direct electrostatically-mediated interaction between the TPR domain and the NTE of MPS1. An MPS1 version designed to bypass this interaction is no longer regulated by Aurora B or itself, and is inefficiently delocalized by microtubules in metaphase. A recent study using chemical crosslinking on recombinant MPS1 showed that crosslinks can be detected between the NTE and TPR, confirming that the modules interact in full-length MPS1 ^58^. We envision a model in which the NTE and TPR transiently interact, preventing efficient binding of MPS1 to kinetochores. The NTE-TPR interaction is diminished by Aurora B activity via unknown mechanisms, thereby enhancing affinity of MPS1 for kinetochores **(Fig S7a)**. Once on kinetochores, the ability of the two modules to interact promotes MPS1 release and enables SAC silencing upon the formation of stable kinetochore-microtubule attachments (**Fig S7b**). The apical NTE which we added in MPS1^2xNTE^ is likely not able to interact with the TPR and is thus always available for interaction with kinetochores, leaving the Aurora B input mute. At the same time, the inability of this extra NTE to interact with TPR reduces MPS1’s release from kinetochores and decreases sensitivity to displacement by microtubules. Of note: since deletion of only the TPR domain reduces MPS1 levels at kinetochores ^31^, this domain has an important contribution in addition to regulation of the NTE, possibly in initial recruitment of the protein. It will be important to define this contribution as well as the mechanisms by which MPS1 is regulated by both MPS1 itself and Aurora B.

The metaphase delay observed in MPS1^2xNTE^-expressing cells is very similar to previously tested conditions in which MPS1 was tethered to kinetochores via fusion to the MIS12 protein ^37^. Consistently, both situations display elevated levels at kinetochores of SAC components downstream of MPS1 activity. Interestingly, however, MPS1^2xNTE^ appears to promote KNL1-MELT phosphorylation and BUB1 recruitment more efficiently than MAD1 recruitment (**Fig 4d,e**, **S6a,b**). In addition, although MAD1 kinetochore levels were low on average in MPS1^2xNTE^ metaphase cells, they were quite variable between kinetochores, implying that only a subset of kinetochores were proficient in generating a strong enough SAC response. MPS1^2xNTE^ may thus expose conditions in which the balance between SAC activating and silencing mechanisms (kinase, phosphatase and dynein) is near a tipping point. As such, MPS1^2xNTE^ may be a useful tool to examine which SAC silencing events are most senstitive to reductions in MPS1 and how.

Previous work showed that microtubules inhibit binding of two regions of MPS1 to two NDC80-C components: a short MR region interacting with NUF2 and the NTE-TPR region interacting with HEC1, the latter in an at least partly non-competitive manner ^32,33^. In support of this, the MPS1^2xNTE^ variant, which is insensitive to the two most prominent regulators of MPS1-NDC80-C interactions (Aurora B and MPS1), was substantially displaced upon microtubule attachments. Surprisingly, however, it didn’t reach the basal levels observed in MPS1^WT^ but was maintained significantly on metaphase kinetochores without noticable effects on microtubule occupancies. This implies that microtubules indeed do not only inhibit MPS1 kinetochore localization through direct disocciation of NTE and NDC80-C but may additionally promote NTE-TPR interactions, for example by the kinetochore enrichment of phosphatases which could promote their actions on the NTE ^32^.

The work presented in this study sheds new light on the complex relationship between MPS1, kinetochores and microtubules and reveals that regulated intramolecular interactions in MPS1 act alongside the microtubule competition to efficiently displace MPS1 from kinetochores. This novel layer of regulation is, therefore, important in ensuring smooth mitotic progression and in enabling SAC silencing once stable end-on microtubule attachments have been formed.

## ACKNOWLEDGEMENTS

We thank all the Kops and Perrakis lab members for suggestions and discussions. We would especially like to acknowledge Prof. Marcellus Ubbink for providing access to the NMR facility at the Leiden Institute of Chemistry (Leiden University) and for critically discussing initial NMR experiments and data, Wouter Touw for help with the TALOS software, and Marvin Tanenbaum and his group for the microscope used for the FRAP imaging and for help with performing those experiments. This work is part of the Oncode Institute, and was funded by grants from the Dutch Cancer Society (KWF/HUBR-2012-5427) and from the Netherlands Organisation for Scientific Research (NWO-Vici 865.12.004).

## MATERIALS AND METHODS

### Isotopically labelled compounds

^15^NH_4_Cl, ^13^C_6_-glucose, D_2_O and ^15^N-asparagine were purchased from CortecNet (Voisins-Le Bretonneux, France)

### Protein production

For the proteins used in the NMR experiments, ^15^N- and ^15^N, ^13^C-enriched minimal media were prepared as described previously^59^. Uniformly isotopically enriched MPS1 samples were produced as following. The plasmids containing the constructs of MPS1 variants were transformed into the BL21(DE3) strain. A single colony was inoculated into 5 mL Lysogeny broth (LB) medium supplemented with 30 μg mL^−1^ kanamycin at 37 °C for 3 hours. 50 μL of the pre-culture was transferred into 50 mL of minimal medium and grown at 37 °C overnight. 10 mL of the pre-culture was then transferred into 1 L of minimal medium and grown at 37 °C until OD600 reached ^~^0.6. Gene expression was induced with 0.5 mM Isopropyl β-D-1-thiogalactopyranoside (IPTG) and the cultures were allowed to grow at 22 °C for 18 hours. Cells were harvested by centrifugation and resuspended in 20 mM KPi, pH 7.5, 1 mM TCEP (buffer A) supplemented with 300 mM KCl, 10 mM imidazole, and 1 mM DNase. Samples were stored at −20 °C before proceeding to purification. The resuspended cells were defrosted at room temperature. The sample was lysed by sonication at 50% amplitude for three minutes with Qsonica Sonicator Q700 (Fisher Scientific). The lysate was further disrupted by EmulsiFlex (Avestin). Following centrifugation at 21,000 g for 20 minutes at 4 °C, the supernatant was loaded on a HisTrap HP column (GE Healthcare). After extensive washing in buffer A supplemented with 500 mM KCl and 5 mM imidazole, the protein was eluted in the same buffer, but now supplemented with 300 mM imidazole. The samples were diluted three-fold in buffer A with 50 mM KCl and loaded on a HiTrap Heparin HP column (GE Healthcare). After washing with the same buffer, the protein was eluted in buffer A containing 500 mM KCl. The sample was then incubated with 3C protease for affinity tag cleavage at 4 °C overnight. The sample was subsequently loaded on a HisTrap HP and HiTrap Heparin column and eluted in buffer A containing 500 mM KCl. The elute was then loaded on a Superdex G75 16/60 HiLoad (GE Healthcare) preequilibrated in 20 mM HEPES/NaOH, pH 7.5, 150 mM KCl, 1 mM TCEP and 7% D_2_O (buffer B). The protein fractions were pooled together and concentrated. The concentration of the MPS1 samples were determined spectrophotometrically using ∊280nm = 9.97 mM^−1^ cm^−1^. The purified proteins were aliquoted and stored at −80 °C. 15N asparagine labeled-labeled MPS1 samples were produced as a protocol adapted from Tong et al^60^.

For the proteins used in the MST experiments, The 1xNTE (WT) and 2xNTE MPS1 constructs (residues 1-377 and 1-808) were cloned into the pFastBac-HT B vector for insect cell expression. Recombinant baculovirus was generated following the manufacturer’s instructions (Invitrogen). Spodoptera frugiperda (Sf9) insect cells were infected with the baculovirus and allowed to grow for 72 hours at 27 °C. Cells were harvested by centrifugation and re-suspended in 50 mL of buffer A supplemented with 150 mM KCl and 10 mM imidazole and one tablet of Pierce™ Protease Inhibitor Tablets EDTA-free (Thermo Fisher Scientific). Samples were stored at - 20°C before proceeding to purification. The re-suspended cells were lysed by sonication for one minute at 50% amplitude in a Qsonica Sonicator Q700 (Fisher Scientific). Following centrifugation at 21,000 g for 20 minutes at 4 °C, the supernatant was incubated with Ni^2+^ charged Chelating Sepharose Fast Flow resin (GE Healthcare) for 30 minutes at 4 °C. After extensive washing in buffer A supplemented with 500 mM KCl and 5 mM imidazole, the protein was eluted in 15 mL of buffer A supplemented with 50 mM KCl and 300 mM imidazole. The eluent containing MPS1 was subsequently diluted two-fold in buffer A with 50 mM KCl and loaded on a HiTrap Q HP column (GE Healthcare) After washing with the same buffer, the protein was eluted in buffer A containing 400 mM KCl. The sample was then loaded on a Superdex G75 16/60 HiLoad (GE Healthcare) pre-equilibrated in 20 mM HEPES/NaOH, pH 7.5, 150 mM KCl and 1 mM TCEP. The protein fractions were pooled together and concentrated to ^~^30 µM. The purified protein stored in 50 µL aliquots, flash-frozen by liquid nitrogen and stored at −80 °C.

### NMR measurements and backbone assignment

The protein concentrations of the ^13^C, ^15^N MPS1 #1-239 and ^13^C, ^15^N MPS1 #62-239 were 450 µM and 550 µM, respectively in buffer B. All NMR spectra were recorded on a Bruker AVIIIHD 850 spectrometer with a TCI-Z-GRAD cryoprobe at 298 K. The 3D HN(CA)CB, HNCA, HN(CA)CO, HNCO and HN(COCA)CB experiments were acquired for the backbone assignment. The data was processed using Topspin 3.1 (Bruker, Biospin) and spectral assignment and analysis was performed using CCPN analysis 2.1.5.^61^. Sequential assignments of the MPS1 TPR-CTE (residues 62-239) and NTE-TPR-CTE (1-239) domains were performed using combination of HN(CA)CB, HNCA, HN(CA)CO, HNCO and HN(COCA)CB. To refine the assignments, the MPS1 samples were selectively labeled with ^15^N asparagine. Finally 209 assignments, 87% of assignable residues, were made for the NTE-TPR-CTE construct. It can be noted that most of the peaks that could not be assigned lie in the region of the CTE motif, presumably due to the intermediate exchange dynamics for NMR timescale.

### Backbone torsion angle restrains calculation

Phi and psi backbone dihedral angle restraints were predicted by TALOS-N^49^ based on chemical shifts of backbone atoms H, N, Cα and Cβ. Only predictions with a majority consensus in the TALOS-N database and predictions that indicated a dynamic conformation were used for modeling and validation.

### Microscale Thermophoresis and analysis

The thermophoresis measurements were performed as previously described (Hiruma et al^62^) with a slight modification. The DY-547P1 labelled NDC80-C Bonsai (ΔHec1 1-80)^63^ was used at a final concentration of 50 nM in the ATP buffer (20 mM HEPES, pH 7.4, 100 mM KCl, 1 mM ATP, 4 mM MgCl_2_, 1mM TCEP, 0.05% Tween20). The measurement was performed in duplicates at 20% LED and 40% MST power. The binding curves were fitted with a standard one site model (equation 1) and two site model (equation 2) using non-linear regression in GraphPad Prism 6 (GraphPad Software, Inc, USA).

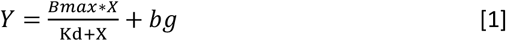

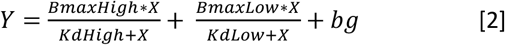

with Y the response; Bmax the maximum response; X the inhibitor concentration; and bg the background response values. An extra sum-of-squares F test was performed with the simpler model being selected unless the p value was under 0.001 to determine which of the two models to select in each case.

### Cell culture and generation of stable cell lines

HeLa Flp-In cells were grown in Dulbecco’s modified Eagle medium (DMEM; Sigma D6429) supplemented with 9% tetracycline-free fetal bovine serum (FBS), penicillin-streptomycin (50 μ g ml^−1^; Sigma P0781) and Ala-Gln (2 mM; Sigma G8541) at 37 °C and 5% CO2. Live-cell imaging was performed in DMEM without phenol red (Sigma; D1145) supplemented with 9% FBS, penicillin-streptomycin (50 μ g ml^−1^; Sigma P0781) and Ala-Gln (2 mM; Sigma G8541). Plasmids were transfected into FlpIn HeLa cells using Fugene HD (Promega) according to the manufacturer’s instructions. To generate stably integrated HeLa Flp-In cell lines, with LAP-tagged genes stably integrated in the FRT site and TetR inducible, pcDNA5 constructs were co-transfected with pOG44 recombinase in a 1:9 ratio and kept in hygromycin (Roche, 10843555001) selection for three weeks.

### Plasmids and cloning

pcDNA5-LAP-MPS1 (WT, Δ60, Δ200) plasmids were generated in ref 31. All other pcDNA5-LAP MPS1 constructs used in this study were generated from the WT plasmid by standard Gibson assembly protocol of PCR products with primers that either amplify the NTE (to generate the 2xNTE) or that contain the desirable point mutations. pcDNA5-LAP-MPS1^N18P-ΔΤPR^ was generated by inserting the N18P mutation with the same strategy but using the pcDNA5-LAP-MPS1^ΔTPR^ that was generated in ref 31 as a template.

### Knockdown, addback and additional cell treatments

For experiments with knockdown-addback of MPS1, siRNAs for GAPDH (as control, Thermo Fisher Scientific; D-001830-01-50, 20nM) or MPS1 (custom, Thermo Fisher Scientific; 5’-GACAGAUGAUUCAGUUGUA-3’, 20nM) were transfected using RNAi Max (Thermo Fisher Scientific) according to manufacturer’s instructions. After 16 h of siRNA treatment, cells were arrested in S-phase by addition of thymidine (2 mM; Sigma-Aldrich cat. no. T1895). To induce expression of exogenous LAP-tagged proteins, doxycycline (1 μ g ml−1; Sigma-Aldrich cat. no.D9891) was added 8 hours after the thymidine addition. After 24h of thymidine addition, cells were released and treated with the indicated drugs: ZM-447439 (2 μ M; Tocris Bioscience, cat. no. 2458); Cpd-5 (250 nM; gift from R. H. Medema, Netherlands Cancer Institute); nocodazole (3.3 μ M; Sigma-Aldrich cat. no. M1404). MG-132 (5 mM; cat. no. C2211) was only added 8-9 hours after thymidine release to first allow cells to enter mitosis. Cells were used for further experiments between 8-10 h after thymidine release. For simultaneous knockdown of HEC1 and MPS1, cells were first transfected with 40 nM siHEC1 (custom; Thermo Fisher Scientific; 5’-CCCUGGGUCGUGUCAGGAA-3’). After 24 hours cells were transfected with a second round of 40 nM siHEC1 and a first round of 20nM siGAPDH or siMPS1. 16 hours later, cells were arrested in S-phase and the protocol continues as above.

### Fixed cell immunofluorescence microscopy and image quantification

For immunofluorescence, HeLa FlpIn cells grown on 12 mm coverslip (no. 1.5) were permeabilized for 1 min with warm PEM buffer (100mM Pipes (pH 6.8), 1mM MgCl2 and 5mM EGTA), followed by fixation for 10 min with 4% PFA in PBS. For analysis of cold-stable microtubules, cells were placed on ice for 15 min prior to pre-extraction and fixation. After fixation, coverslips were washed three times with PBS and blocked with 3% BSA in PBS overnight at 4°C. The next day, primary antibodies diluted in 3% BSA were added to the coverslips and incubated for 2 h at room temperature. Subsequently, cells were washed three times with 0.1% triton in PBS and incubated with secondary antibodies in 3% BSA for another hour at RT. Coverslips were then washed two times with 0.1% Triton in PBS followed by 2 min incubation with DAPI diluted in PBS, followed by two final washes in PBS. Coverslips were then mounted onto glass slides using Prolong Gold antifade. All images were acquired on a deconvolution system (DeltaVision Elite Applied Precision/GE Healthcare) with a × 100/1.40 NA UPlanSApo objective (Olympus) using SoftWorx 6.0 software (Applied Precision/GE Healthcare). Images were acquired as z-stacks at 0.2 μ m intervals and all images of similarly stained experiments were acquired with identical illumination settings. Images were then deconvolved and maximum intensity projections were made using SoftWoRx. Cells were selected based on the mitotic shape of DAPI signal.

For quantification of images, a CellProfiler^64^ pipeline was used to threshold and select all kinetochores and all chromosome areas (excluding kinetochores) using the DAPI and CENP-C. This was used to calculate the relative average kinetochore intensity of various proteins ((kinetochores - chromosome arm intensity (kinetochore localized protein of interest))/(kinetochores-chromosome arm intensity (CENP-C))).

### Live cell imaging and movie analysis

For simple live-cell imaging of mitotic timing in various conditions, cells were plated in 24-well plates and DIC filming started 6 hours after thymidine release on a Nikon Ti-E motorized microscope equipped with a Zyla 4.2Mpx sCMOS camera (Andor) and 40× 1.3 NA objective lens (Nikon). Cells were kept at 37 °C and 5% CO2 using a cage incubator and Boldline temperature/CO2 controller (OKO-Lab). In experiments with DNA visualization, SiR-DNA (Spirochrome, 100nM) was added right after thymidine release and images acquired were comprised of 8 z-slices separated by 2μm with the same timing as above. Fluorescence excitation was done using Spectra X LED illumination system (Lumencor) and Chroma-ET filtersets.

Analysis of live-cell imaging experiments was carried out with ImageJ software. Time in mitosis for DIC movies was defined as the time between nuclear envelope breakdown (defined as cell rounding) and anaphase-onset or cell flattening. For movies where DNA was visualized, mitosis onset was defined as the first frame where chromatin condensation was observed.

### Fluorescence recovery after photobleaching (FRAP)

Cells were grown in 96-square-well glass bottom dishes (Matriplate, Brooks). They were subsequently treated with siMPS1 and expression of MPS1^WT^ or MPS1^2xNTE^ was induced. Cells were arrested in S phase with thymidine and after 24 hours released in nocodazole. 30 minutes before imaging, Cpd-5 was added to inactivate MPS1 and exclude activity dependent changes in turnover and MG132 to prevent cells from exiting mitosis. Mitotic cells were selected and images were acquired using a Yokogawa CSU-X1 spinning disk confocal attached to an inverted Nikon TI microscope with Nikon Perfect Focus system, 100× NA 1.49 objective, an Andor iXon Ultra 897 EM-CCD camera, and Micro-Manager^65^ and NovaLum software. The EYFP-based LAP tag of LAP-MPS1 was bleached using the 488-nM laser line (Andor FRAP laser box) set to 100%. Areas centered on single kinetochore pairs were bleached once at 100% laser power for 2,000 ms. Fluorescence intensity of the entire cell was acquired for 20 pre-bleach iterations and for 90 iterations after bleach with 100ms intervals.

For each time point, the mean fluorescence intensity was measured in the area that encompassed kinetochore movement and in a similarly sized directly neighboring cytosolic area that was devoid of kinetochores throughout the experiment. Both areas were corrected for background, and the mean fluorescence of the cytosolic area was subtracted from the kinetochore area for each time point (area_(KT-cyto)_). For each measurement, the fluorescence intensity of the area_(KT-cyto)_ in the timepoint before bleaching was set to 100%, and the measured postbleach area_(KT-cyto)_ signal was normalized to this value. The signal from the bleached kinetochores was corrected for general cell-wise loss in fluorescence by measuring the decline in signal in a non-bleached kinetochore pair. Recovery half-times were determined by nonlinear curve fitting based on a one-phase association followed by a plateau using Prism software (GraphPad Software).

### Antibodies

The following primary antibodies were used: CENP-C (polyclonal guinea pig, 1:2,000; MBL, Catalog#: PD 030), a-Tubulin (mouse monoclonal, 1:10,000; Sigma-Aldrich, Catalog#: T5168), HEC1 (mouse monoclonal 9G3, 1:500; Thermo Fisher Scientific, Catalog#: MA1-23308), GFP (custom rabbit polyclonal raised against full-length GFP as antigen, 1:10,000), GFP (mouse monoclonal, 1:1,000; Sigma, Catalog#: 11814460001), MAD1 (mouse monoclonal, 1:1000; Merck Millipore, Catalog#: MABE867), BUB1 (rabbit polyclonal, 1:1,000; Bethyl, Catalog#: A300-373 A-1), Astrin (rabbit polyclonal, 1:1000; Bethyl, Catalog#: A301-511A), MPS1-NT (mouse monoclonal, 1:1000; EMD Millipore, Catalog#: 05-682), KNL1-pT180 (custom rabbit serum against pT180). Secondary antibodies (Invitrogen Molecular Probes, all used at 1:600) were goat anti-guinea pig Alexa Fluor 647 (Catalog#: A21450), goat anti-rabbit Alexa Fluor 488 (Catalog#: A11034), 568 (Catalog#: A11036), and anti-mouse Alexa Fluor 488 (Catalog#: A11029) and 568 (Catalog#: A11031).

### Statistics and reproducibility

Results from immunofluorescence images from different experiments were pooled and no difference was observed between different experimental sets. The comparisons most pertinent for the conclusions and number of independent experiments are specified in the figures and legends. Two-tailed, unpaired t-tests or one-way ANOVA followed by Tukey’s test were performed to compare either two or multiple experimental groups respectively in immunofluorescence quantifications when n ≥ 3. In those cases in which n < 3, no statistical analysis was performed. Data are presented as mean ± s.d. All replicates showed similar results and a representative experiment was displayed.

---

**Figure S1.**
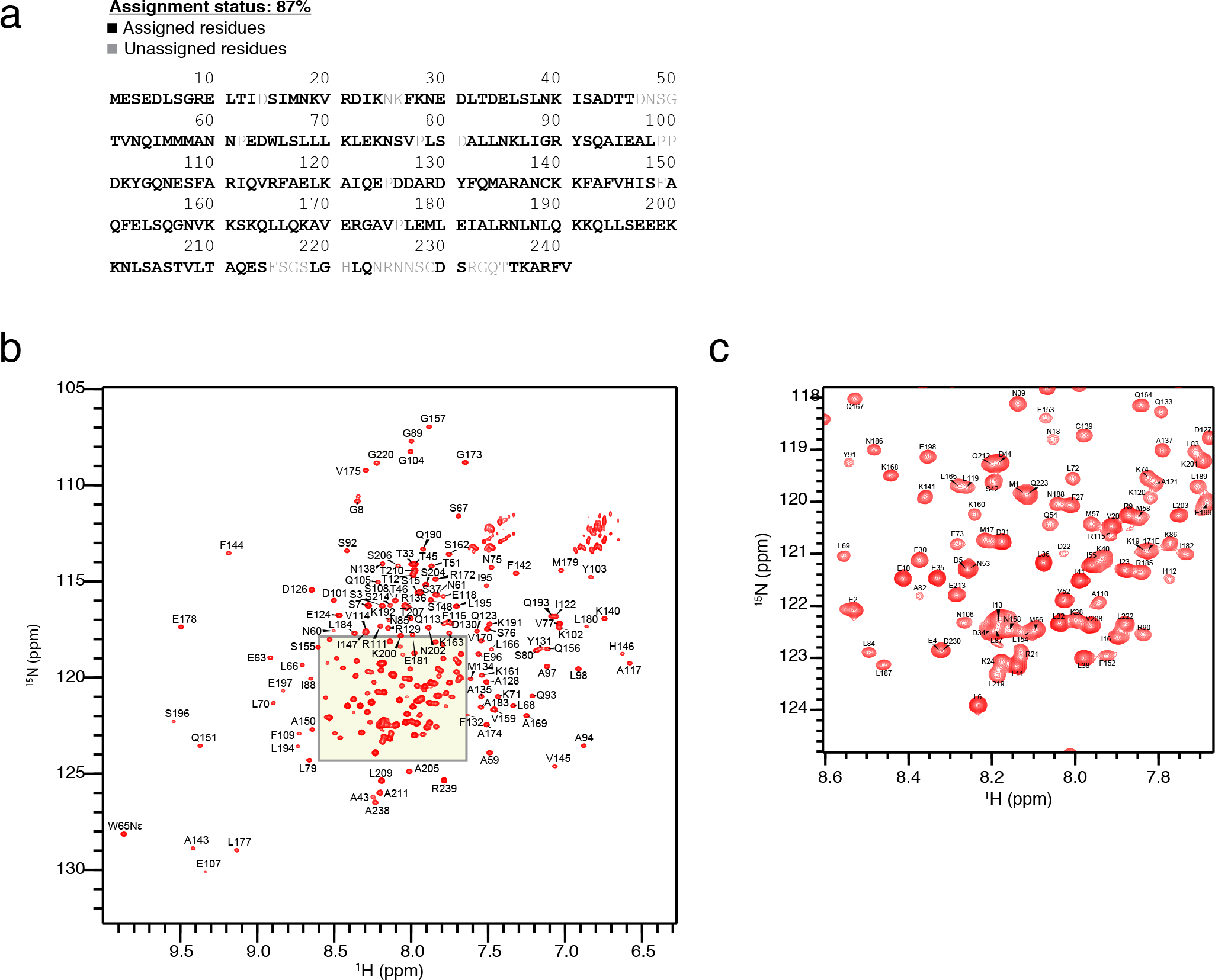
NMR residue assignment. **(a)** Assignment status after NMR of the MPS1 N-terminal region displayed on the amino-acid sequence. NMR assigned residues annotated in black, unassigned in grey. **(b and c)** NMR spectra of MPS1 N-terminal region and residue assignment **(b)** with zoom in on cluster of spectra **(c)**.

**Figure S2.**
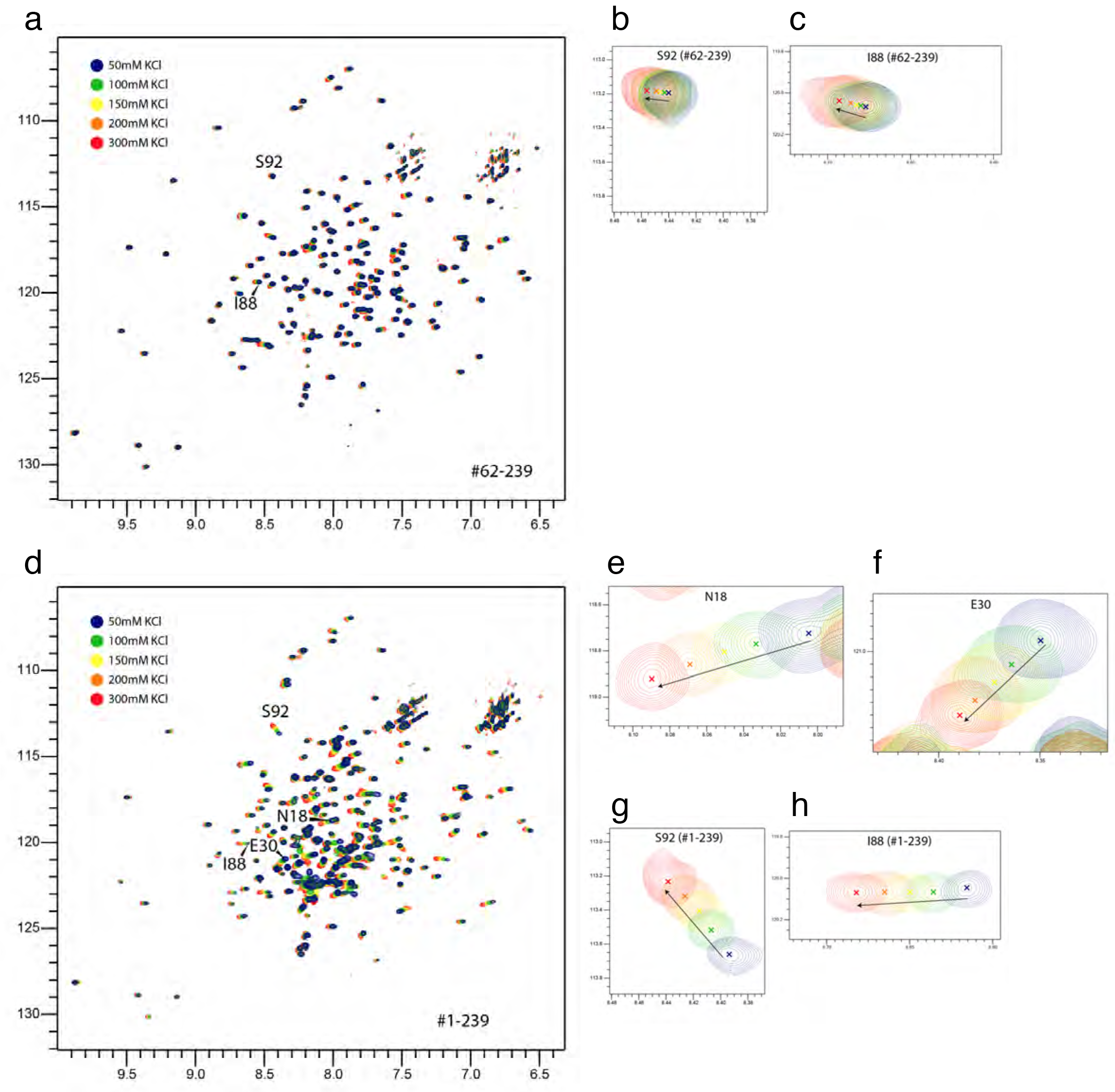
NMR of MPS1 upon KCl titration. **(a-c)** NMR spectra of MPS1^62-239^ **(a)** and examples of individual residue spectra **(b,c)** in different KCl concentrations. **(d-h)** NMR of MPS1^1-239^ **(d)** and examples of individual residue spectra **(e-h)** in different KCl concentrations.

**Figure S3.**
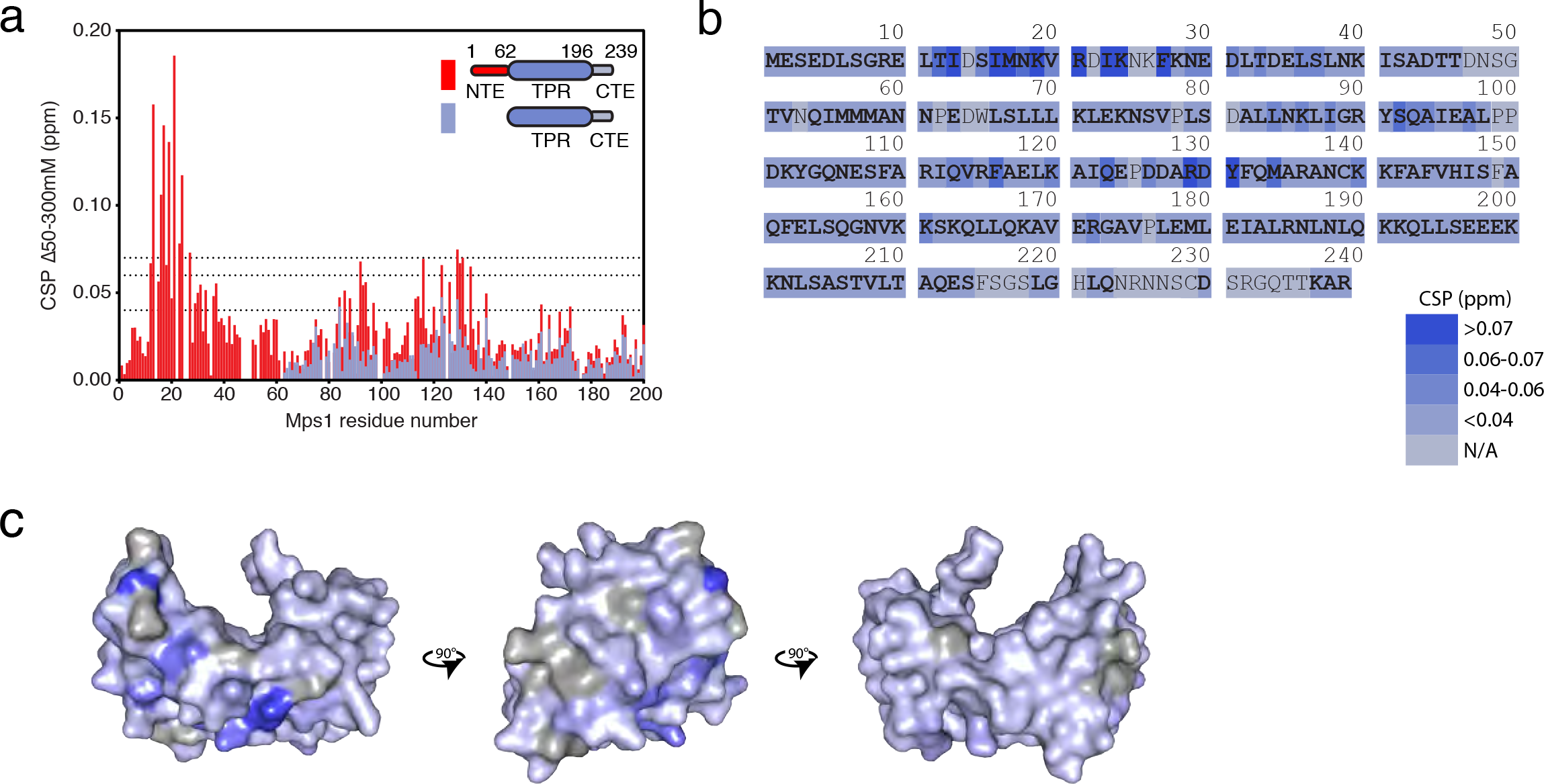
The NTE-TPR interaction is electrostatic. **(a)** Quantification of the chemical shift perturbations (CSPs) per residue in the NTE-TPR-CTE region between the lowest and highest KCl concentrations, colour-coded based on the MPS1 fragment used. **(b)** Display of the CSPs onto the sequence of the N-terminal region per residue, colour-coded based on the CSP magnitude. **(c)** Projection of the CSPs onto the crystal structure of the TPR domain in 3 orientations, following the same colour-coding as in (b).

**Figure S4.**
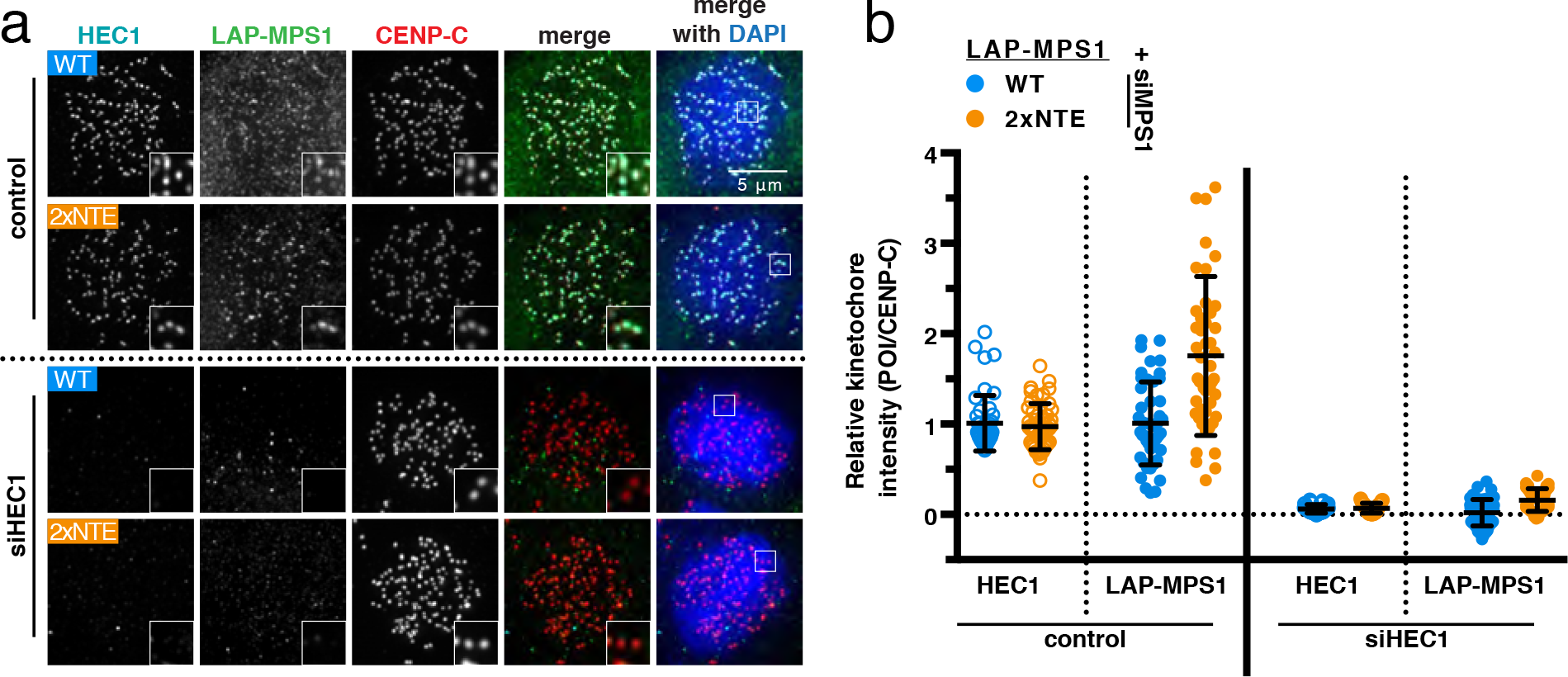
MPS1^2xNTE^ kinetochore localization depends on NDC80C. **(a and b)** Representative images **(a)** and quantification **(b)** of the kinetochore levels of the indicated proteins in nocodazole-treated HeLa Flp-In cells transfected with MPS1 and GAPDH (control) siRNA or with MPS1 and HEC1 siRNA and expressing the indicated LAP-MPS1 variants The graph displays the mean kinetochore intensity (±s.d.) normalized to the levels of each protein in MPS1^WT^-expressing cells treated with control siRNA. Each dot represents one cell and the experiment was performed two times.

**Figure S5.**
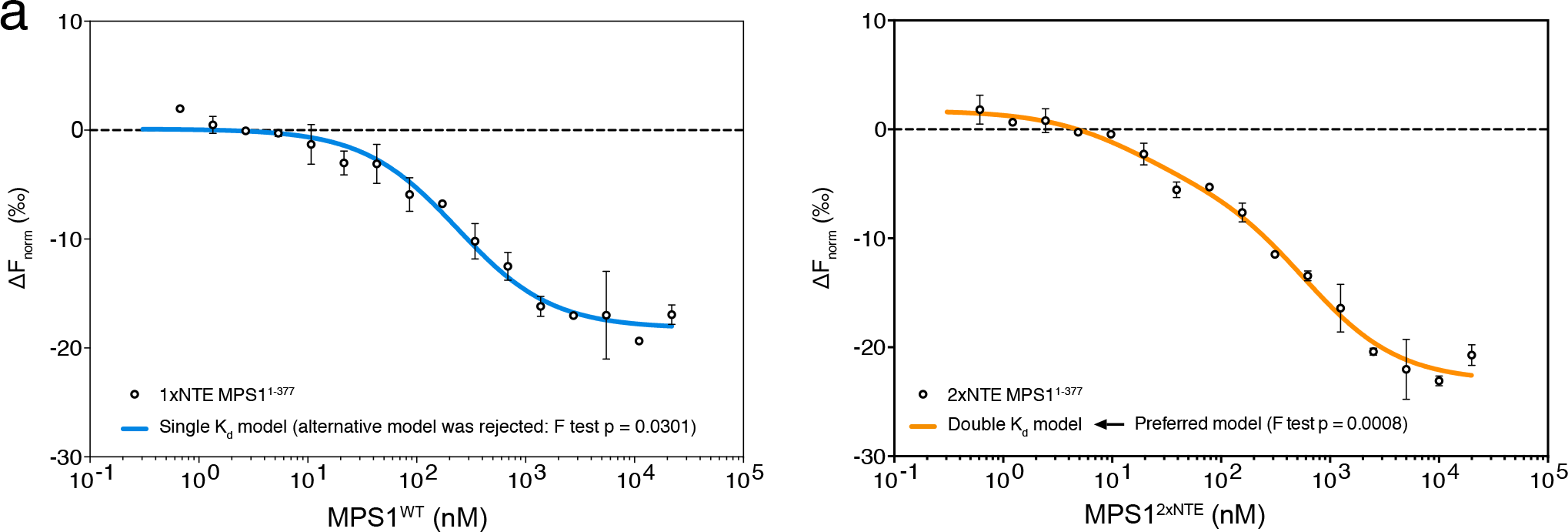
MST of MPS1^1-377^. **(a)** MST binding graph generated by titrating MPS1^WT^ (left panel) or MPS1^2xNTE^ (right panel) to 50nM of NDC80^Bonsai^. One- or two-site binding curves were fitted and F test was performed to select a preferred model.

**Figure S6.**
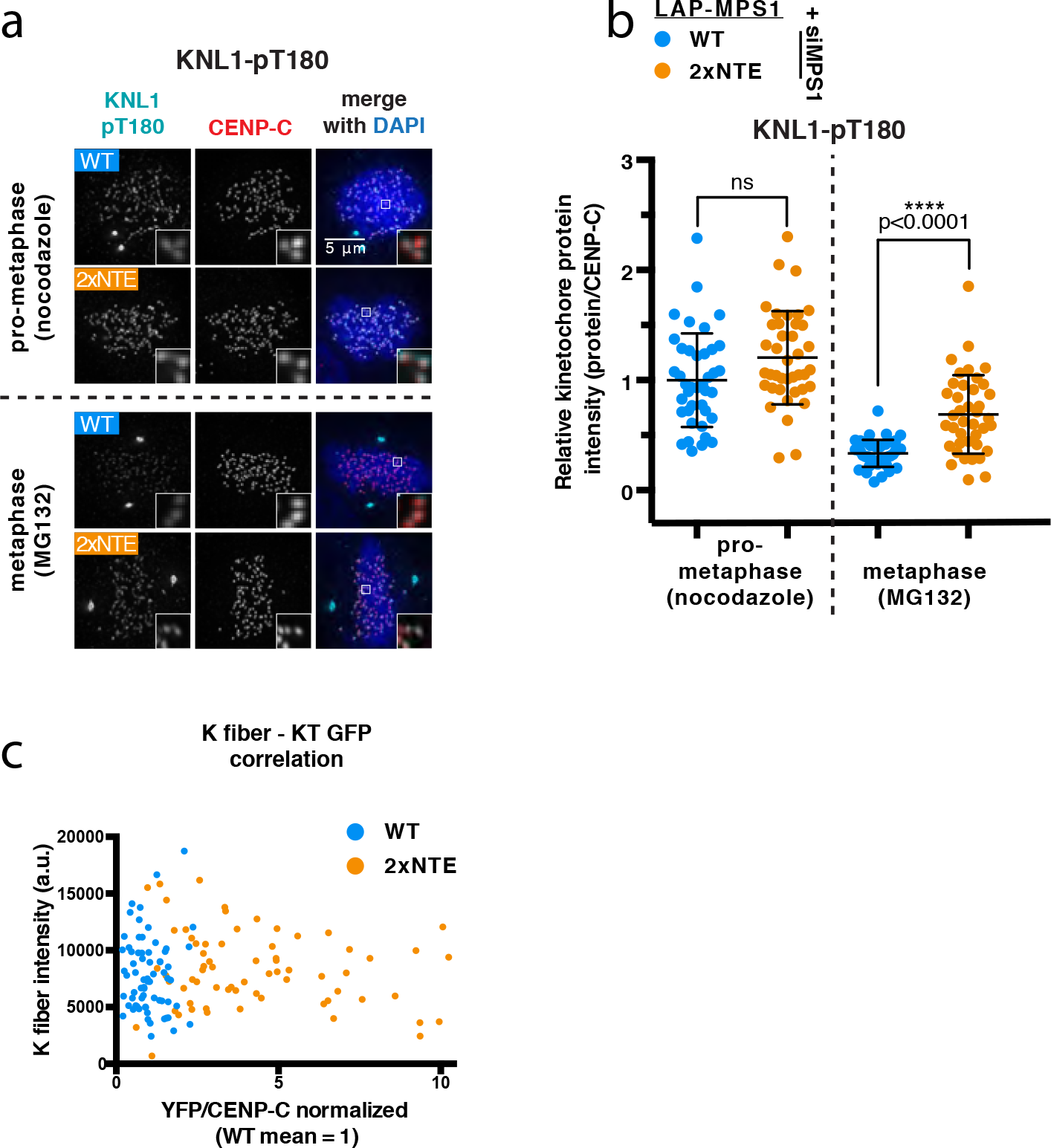
High levels of MPS1^2xNTE^ on kinetochores causes substrate phosphorylation without preventing microtubule binding. **(a and b)** Representative images **(a)** and quantification **(b)** of the kinetochore levels of KNL1-pT180 in nocodazole- or MG132-treated HeLa Flp-In cells transfected with MPS1 siRNA and expressing the indicated LAP-MPS1 variants. The graph displays the mean kinetochore intensity (±s.d.) normalized to the levels of KNL1-pT180 in prometaphase MPS1^WT^-expressing cells. Cells were pooled from three independent experiments and each dot represents one cell. Asterisks indicate significance (student’s t test was performed between the cell lines in each condition). **(c)** Correlation plot of the kinetochore levels of LAP-MPS1 with k-fiber intensity on individual kinetochores, from HeLa Flp-In cells transfected with MPS1 siRNA and expressing the indicated LAP-MPS1 variants. LAP-MPS1 kinetochore levels were normalized to the levels of MPS1^WT^. Each dot represents one kinetochore and measurements were pooled from three independent expriments.

**Figure S7.**
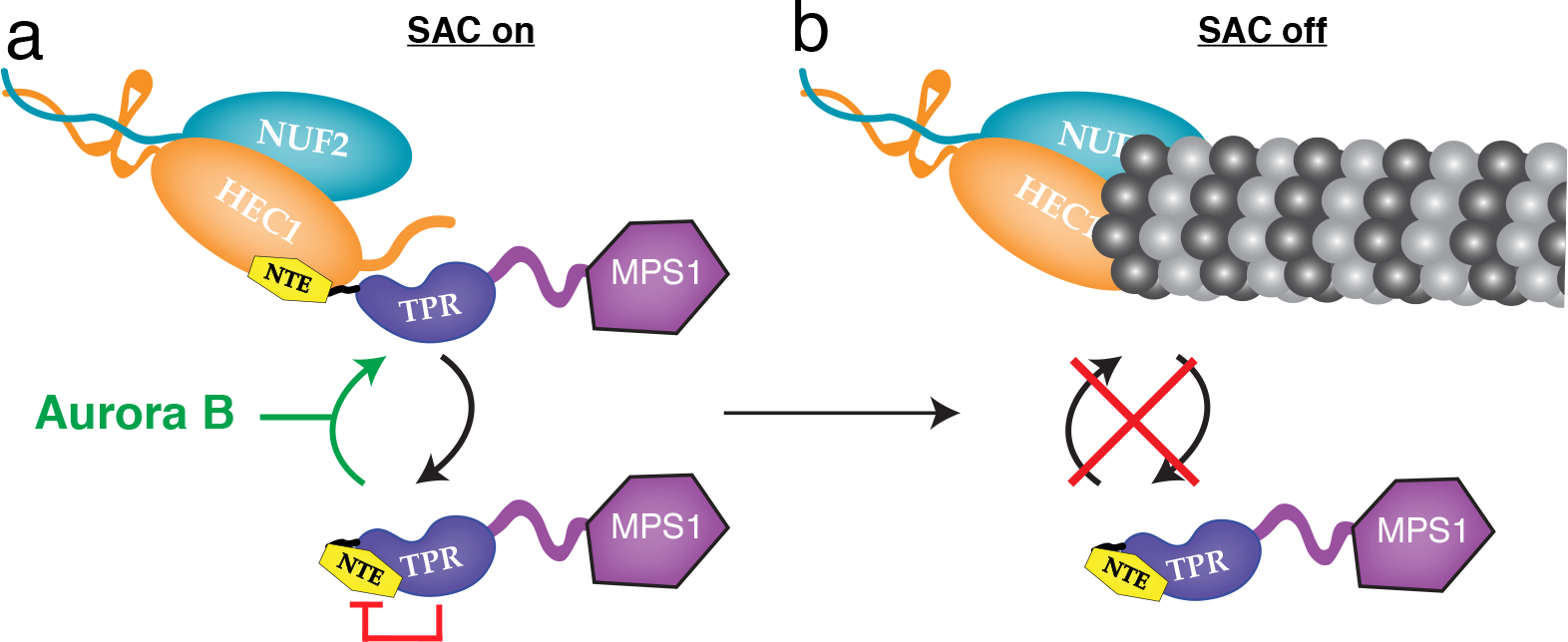
Model for regulation of kinetochore localization of MPS1 via an NTE-TPR interaction. In MPS1^WT^ the NTE and TPR transiently interact, preventing efficient binding of MPS1 to kinetochores. The NTE-TPR interaction is diminished by Aurora B acitivity, allowing MPS1 to bind kinetochores **(a)**. Once on kinetochores, the ability of the two modules to interact promotes MPS1 release and enables SAC silencing to occur upon the formation of stable kinetochore-microtubule attachments **(b)**.

